# Inter-laboratory harmonization of microsphere immunoassays for SARS-CoV-2 antibody detection in dried blood spots and oral fluids

**DOI:** 10.1101/2024.10.19.619238

**Authors:** Kate L. DeRosa, Nora Pisanic, Kate Kruczynski, Christopher D. Heaney, Linda M. Styer, Nicholas J. Mantis

## Abstract

Dried blood spots (DBS) and oral fluids (OF) are easily attainable biospecimen types that have enabled population scale antibody monitoring for SARS-CoV-2 exposure and vaccination. However, the degree to which the two different biospecimen types can be used interchangeably remains unclear. To begin to address this question, we generated contrived DBS (cDBS) and OF (cOF) from serum panels from SARS-CoV-2 infected, vaccinated, and uninfected individuals. The contrived samples were evaluated using SARS-CoV-2 multiplexed microsphere immunoassays (MIAs) at two different institutions. Intra-laboratory tests revealed near perfect agreement between cDBS and cOF for N and S antigens, as evidenced by κ = 0.97-1 and 98%-100% agreement. Inter-laboratory comparisons were equally robust for both N (κ = 0.94-0.96; 97.5%-98 % agreement) and S (κ = 0.98 -1.0; 99.0%-100%). Furthermore, assays were transferred between labs, including methods and reagents, and a subset of cDBS and cOF samples (n = 52) were tested. Qualitative concordance remained high (κ = 0.94-1.0; 97.5%-100% agreement), confirming that integrity of the assays is retained upon transfer. In summary, our results provide evidence that DBS and OF can be used interchangeably across laboratories and institutions for the qualitative assessment of SARS-CoV-2 antibody determinations.

## 1. Introduction

COVID-19 continues to persist globally due to the extremely high transmissibility rates of SARS-CoV-2 and its ever-evolving variants of concern (VoC). From the standpoint of public health, real time estimates of the vulnerability of distinct cohorts across all age groups to circulating SARS-CoV-2 variants is of paramount importance when making decisions about next generation vaccine implementation. While serological screening can be conducted as a routine part of health care visits, such an approach will be of limited scope and bias towards specific subpopulations. An alternative is to perform serosurveys in the field using low-cost methods that are conducive to self-collection and biospecimen retrieval, then sending samples to one or more centralized testing sites for analysis of antibody and functionality against SARS-CoV-2 VoCs.

Dried blood spots (DBS) and oral fluids (OF) are two biospecimen types that are amenable to self-collection on a cohort, community and even population scale ^1-6^. DBS are not only amenable to self-collection with preassembled kits and simple instructions, but the resulting spots are stable on Whatman filter paper for long periods of time under refrigeration (4-8°C) or frozen (up to -20°C) in the presence of desiccants ^4,7^. Similarly, OF represent an easily accessible biospecimen type to interrogate both systemic and mucosal immune responses even among children ^3^. Thus, DBS and OF are complementary in the sense that they are convenient to collect and amenable to population wide surveys. However, the degree to which the saliva and DBS are interchangeable in terms of SARS-CoV-2 antibody responses has not been established. In this study, we performed a comprehensive side-by-side comparison of paired contrived DBS (cDBS) and oral fluid (cOF) specimens using two SARS-CoV-2 multiplex immunoassays (MIAs) to evaluate the comparative strengths and weaknesses of each sample type.

## 2. Materials and Methods

### 2.1 MIA

In Lab A: Antigens and controls were coupled covalently to magnetic microparticles (hereafter beads, Luminex xMAP) using carbodiimide coupling chemistry with Sulfo-N-hydroxysuccinimide (Sulfo-NHS) as described previously (5 μg protein/1×10^6^ beads, see Table S1 for list of antigens). The assay included a control bead indicating non-specific binding of sample to beads. After activation with the crosslinker, assay buffer (PBS-TBN, Luminex, TX) containing 1 mg/mL BSA was added to this bead set. Each well contained 1,000 beads per set in assay buffer. Detailed assay conditions are provided in Table 1. Assay plates included A blank (assay buffer) on each plate. Plates were read on a MagPix instrument. The blank was subtracted to calculate net median fluorescence intensity (MFI), followed by subtraction of the within-sample background (BSA bead net MFI). Cutoffs were defined based on receiver-operator characteristics (ROC) area under the curve (AUC) for each antigen. Samples above the cutoff of nucleocapsid (GenScript N) and spike (Mt. Sinai S) were classified as ‘prior infection’; samples negative for N but positive for S were classified as ‘prior vaccination’. For cOF, the RBD result was used to classify samples instead of S. Samples below the S/RBD cutoff were classified as naïve, irrespective of the N IgG result.

**Table 1.**
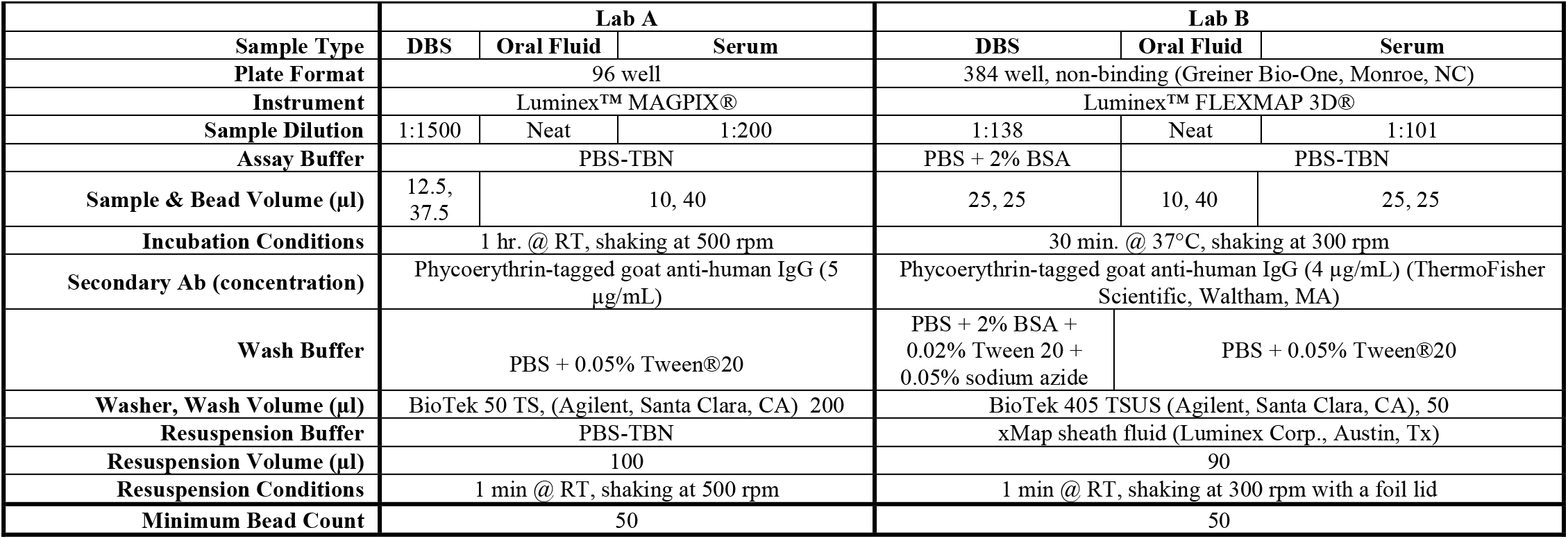
Optimized assay conditions for Lab A and Lab B SARS-CoV-2 MIAs.

In Lab B, SARS-CoV-2 and control antigens listed in **Table S1** were covalently coupled to Magplex-C microspheres (1 ×10^6^ per mL) as described ^1^. Bead mixes were prepared using 1.25 µl of each bead and 13.75 µl of assay buffer (1,250 beads/bead set/well) for cDBS and serum. For cOF, the bead mix was prepared using 2 µl of each bead and 22 µl of assay buffer (2,000 beads/bead set/well). cDBS were eluted and tested as described ^4^. Sera were diluted 1:101 using PBS-TBN in a round-bottom, nontreated, polystyrene 96-well plate (Corning, Corning, NY). cOF (neat) were aliquoted into a round-bottom, nontreated, polystyrene 96-well plate (Corning). Samples and beads were incubated following the protocol outlined in **Table 1**. Samples were washed twice, incubated with secondary antibody, and washed twice more (**Table 1**). Samples were resuspended (**Table 1**) and analyzed using a FLEXMAP 3D® instrument (Luminex Corp., Austin, TX). Results are reported as median fluorescent intensity (MFI) for each bead set. Cutoff values were set for each antigen based on the analysis of 87 pre-COVID pandemic sera (Access Biologicals, Vista, CA). The reactive cutoff was mean MFI +6 standard deviations for S and N antigens for cDBS and S antigens for cOF. The reactive cutoff was mean MFI +3 standard deviations for N antigens in serum and cOF (**Table S2**). N antigens have a higher level of non-specific binding in naµve sera and cOF, increasing the standard deviation, resulting in an artificially inflated reactive cutoff when the +6 standard deviation cutoff is used (**Figure S1**). Specimens reactive for ≥ 2 of the 4 spike antigens (RBD, S1, FLS, TRI) are classified as Spike reactive. Index values were calculated by dividing the sample MFI by the reactive cutoff for each bead set. Index values and log_10_ index values were used for plotting. An index value >1 equivalent to a log index > 0 are considered reactive.

### 2.2 Preparation of contrived DBS and OFs

Contrived specimens were prepared using commercially available serum panels and one confirmed Delta positive specimen (**Table S3**). Freshly collected Type O EDTA whole blood (ZenBio Inc., Durham, NC) was centrifuged for 8 min at 2200 RCF and plasma removed. Because most commercially available whole blood now contains SARS-CoV-2 antibodies, the blood cells were transferred to a sterile tube and triple washed by performing the following three times: adding an equal volume of PBS as cells, mixing by inversion for 10 min and centrifuging for 8 minutes at 2200 RCF. cDBS were then prepared as previously described ^1^. cOFs were prepared using Utak® SMx Oral Fluid (Utak, Valencia, CA) diluted 1:8 using PBS-TBN. Serum was diluted 1:200 in 1:8 SMx Oral Fluid.

### 2.3 Antibody determinations in cDBS and cOFs

Specimens were tested following the above-described procedures by Labs A and B (**Table 1**). In addition, labs A and B exchanged methods and reagents and tested a subset (n=52) of the contrived specimens. There are two aspects of lab A and B’s MIAs that are the same – both use Luminex™ instrumentation and both test plasma/serum (**Table 1**). Aside from these components, the two assays differ in their plate format, sample dilutions and bead mix to sample ratios, and buffers (**Table 1**). Method transfer was performed by both laboratories with minimal changes from the originating lab’s protocol, however, both labs kept their respective wash volumes and plate formats while performing the other laboratory’s protocol. In addition, lab B performed lab A’s assay using Luminex™ FLEXMAP 3D® instrumentation due to lack of access to a Luminex™ MAGPIX®. Lab A did change Luminex™ instruments to a FLEXMAP 3D® while running lab B’s assay to match their protocol.

### 2.4 Data Analysis

For Lab A, index and log index values for Gen N, Mt. Sinai S, and Sino RBD were used for plotting. An index value > 1, or a log index value > 0, is reactive. BSA subtracted net MFIs and cutoffs were adjusted for log transformation by replacing negative values with 1. Lab B’s reactivity was determined for the five SARS-CoV-2 antigens (N NA, RBD, S1, FLS, TRI) using the above-described methods. N and NHT were excluded from analysis due to poor performance in serum and cOF (**Figure S1**).

Qualitative intra- and inter-laboratory concordance was assessed using percent agreement, Cohen’s kappa, and Fleiss’ kappa as calculated using R package irr (v0.84.1; Gamer et al., 2019). Quantitative intra- and inter-laboratory concordance was assessed using Bland Altman plots with a threshold of 95% of data points falling within the 95% confidence interval to be considered concordant (Giavarina, 2015). All analyses and figures were performed in R (v4.3.0; R Core Team, 2023). All figures were created using ggplot2 (v3.4.2; Wickham, 2016) and ggpubr (v0.6.0; Kassambara, 2023).

Specimen reactivity from method transfer between labs A and B was assigned as described above.

## 3. Results

### 3.1 Intra-laboratory cDBS and cOF concordance

To assess intra-laboratory concordance between sample types, a total of 197 serum samples were used to generate paired cDBS and cOF **(Table S3**). First, we assessed the within-laboratory concordance between cDBS and cOF for Lab A’s MIA.. Kappa coefficients of 1 (100% agreement) for both N and S reactivities in this MIA were achieved, indicating perfect qualitative agreement between cDBS and cOF (**Table 2**). N and S reactivities also aligned with the known SARS-CoV-2 status (κ = 0.97-0.99; 97.5%-99% agreement) of donor serum samples. Thus, qualitative concordance between cDBS and cOF was achieved using the Lab A MIA. Quantitative concordance was also achieved for N and S reactive samples based on Bland Altman analysis ^8^ (100.0%, 97.9% within the 95% CI, respectively) (**Figure 1, Table S4**).

**Table 2.**
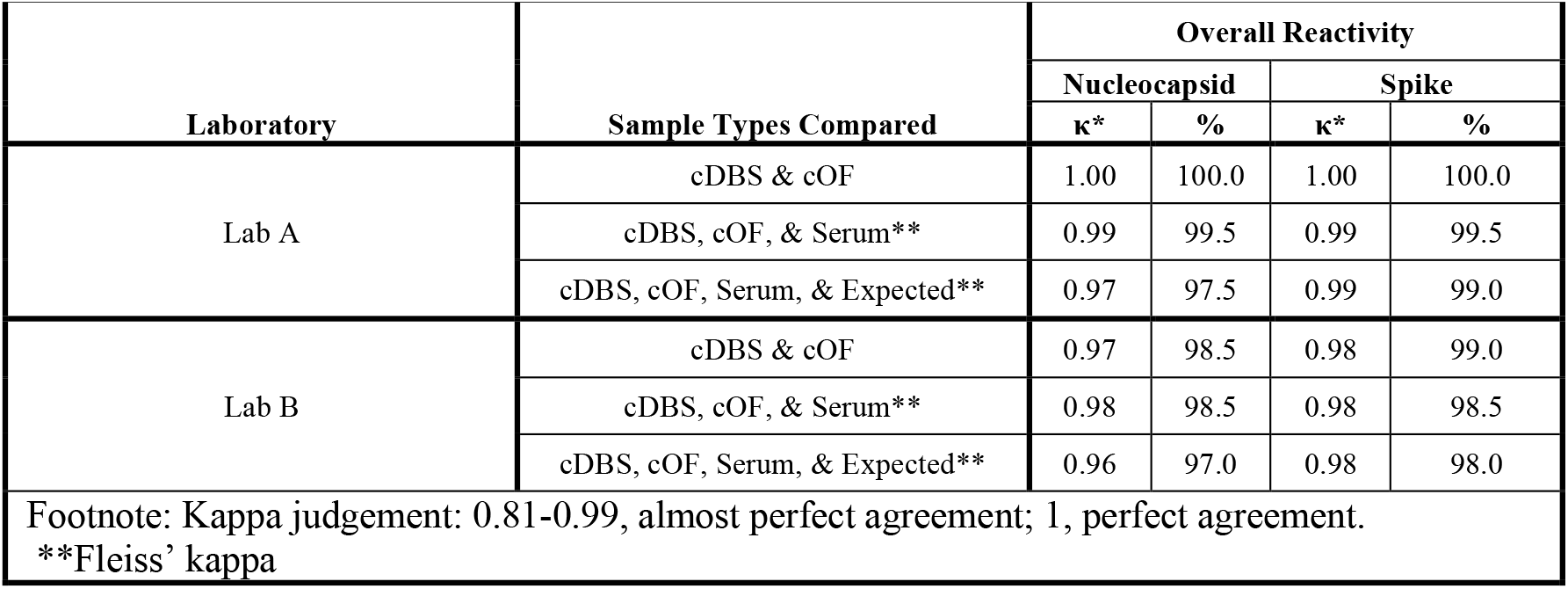
Intra-laboratory kappa coefficient and percent agreement between sample types and expected results for the lab A and lab B MIAs (^*^p = 0).

|  |  |  |  |  |  |
| --- | --- | --- | --- | --- | --- |
| **Laboratory** | **Sample Types Compared** | **Overall Reactivity** |  |  |  |
|  |  | **Nucleocapsid** |  | **Spike** |  |
|  |  | **κ\*** | **%** | **κ\*** | **%** |
| Lab A | cDBS & cOF | 1.00 | 100.0 | 1.00 | 100.0 |
|  | cDBS, cOF, & Serum** | 0.99 | 99.5 | 0.99 | 99.5 |
|  | cDBS, cOF, Serum, & Expected** | 0.97 | 97.5 | 0.99 | 99.0 |
| Lab B | cDBS & cOF | 0.97 | 98.5 | 0.98 | 99.0 |
|  | cDBS, cOF, & Serum** | 0.98 | 98.5 | 0.98 | 98.5 |
|  | cDBS, cOF, Serum, & Expected** | 0.96 | 97.0 | 0.98 | 98.0 |
| Footnote: Kappa judgement: 0.81-0.99, almost perfect agreement; 1, perfect agreement.<br>**Fleiss' kappa |  |  |  |  |  |

**Figure 1:**
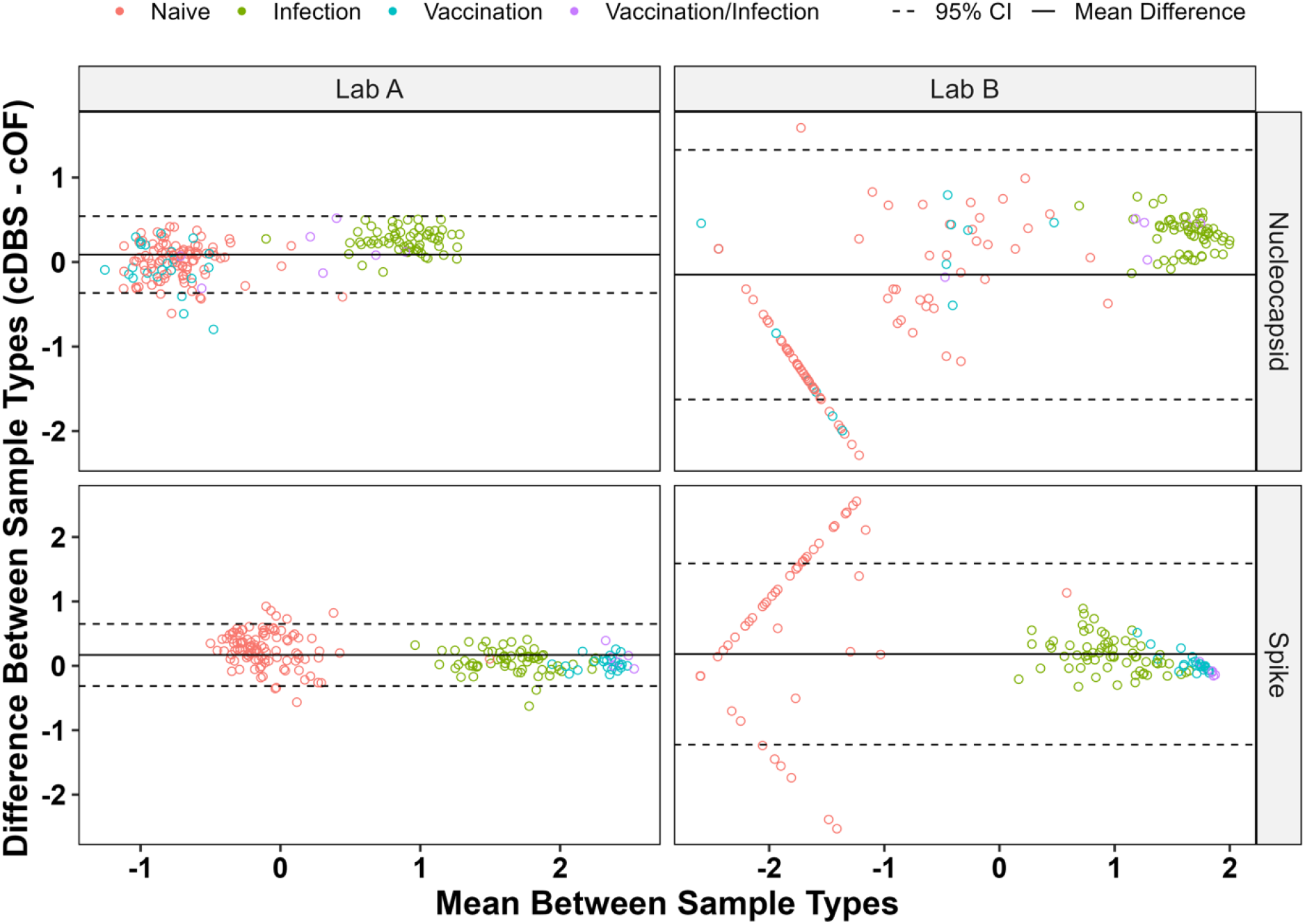
Intra-Laboratory Bland-Altman Analysis for the Lab A and B Assays. Bland Altman plot of 197 paired samples, with comparisons made between cDBS and cOF for the lab A and lab B MIAs. Known SARS-CoV-2 antibody status is shown

Lab B also achieved qualitative and quantitative concordance between cDBS and cOF, but two modifications to the original 8-plex MIA were required. First, the BSA-containing buffers used in that MIA proved to be incompatible with cOF. As a result, Lab B adopted Lab A’s buffers for testing cOF specimens (**Table S5**). Second, the N and NHT antigens in the 8-plex MIA exhibited an unusually high level of non-specific binding in cOF, resulting in elevated cutoff values and false negative classifications (**Figure S1**). Removal of the N and NHT antigens improved assay sensitivity (**Figure S1**). Consequently, qualitative concordance between cDBS and cOF in lab B’s MIA assay was high, based on kappa coefficients for N (κ = 0.97, 98.5% agreement) and S (κ = 0.98, 99% agreement) (**Table 2**). N and S reactivities also agreed with known SARS-CoV-2 status (κ = 0.96 and 0.98; 97%, and 98% agreement, respectively).

Bland Altman analysis ^8^ indicates the lab B assay produces quantitively concordant results for both N and S reactive cDBS and cOF (both 100% within 95% CI) (**Figure 1** (**Table S4**).

### 3.2 Inter-Laboratory cDBS and cOF concordance

Next, we assessed qualitative concordance between lab A and lab B MIA results for cDBS and cOF. Kappa coefficients indicate perfect qualitative agreement for N and S in both sample types (κ = 1.0, > 98% agreement between Lab A and Lab B’s cDBS and cOF results. Moreover, those classifications matched known SARS-CoV-2 status (κ = 0.96-0.99; 95.4%-98.5% agreement) (**Table 3**). Furthermore, cDBS and cOF positive samples were quantitatively concordant for N and S antigens 100.0% of positive samples within the 95% CI), as determined by Bland Altman analysis (**Figure 2**) (**Table S6**). In conclusion, labs A and B successfully demonstrated inter-laboratory qualitative and quantitative concordance for cDBS and cOF.

**Table 3.** Kappa coefficient and percent agreement across sample types and expected results for the lab A and lab B MIAs (^*^p = 0).

|  |  |  |  |  |  |  |
| --- | --- | --- | --- | --- | --- | --- |
| Lab(s) | MIA Method Performed | Sample Types Compared | Overall Reactivity |  |  |  |
|  |  |  | Nucleocapsid |  | Spike |  |
|  |  |  | κ* | % | κ* | % |
| Labs A & B | Labs A & B, respectively | cDBS | 1.0 | 98.0 | 1.0 | 99.0 |
|  |  | Serum | 0.94 | 97.5 | 0.99 | 99.5 |
|  |  | cOF | 0.94 | 97.5 | 1.0 | 100.0 |
|  |  | cDBS, cOF, Serum** | 0.96 | 96.5 | 0.99 | 98.5 |
|  |  | cDBS, cOF, Serum, Expected** | 0.96 | 95.4 | 0.99 | 98.0 |
| Labs A & B | Lab A | cDBS | 1.0 | 100.0 | 0.96 | 98.1 |
|  |  | Serum | 1.0 | 100.0 | 0.96 | 98.1 |
|  |  | cOF | 0.96 | 98.1 | 0.92 | 96.2 |
|  |  | cDBS, cOF, Serum** | 0.99 | 98.1 | 0.96 | 96.2 |
| Labs A & B | Lab B | cDBS | 1.0 | 100.0 | 1.0 | 100.0 |
|  |  | Serum | 0.92 | 96.2 | 0.96 | 98.1 |
|  |  | cOF | 0.61 | 82.7 | 0.96 | 98.1 |
|  |  | cDBS, cOF, Serum** | 0.86 | 82.7 | 0.97 | 96.2 |
| Footnote: Kappa judgement: 0.81-0.99, almost perfect agreement; 1, perfect agreement. **Fleiss' kappa |  |  |  |  |  |  |

**Figure 2:**
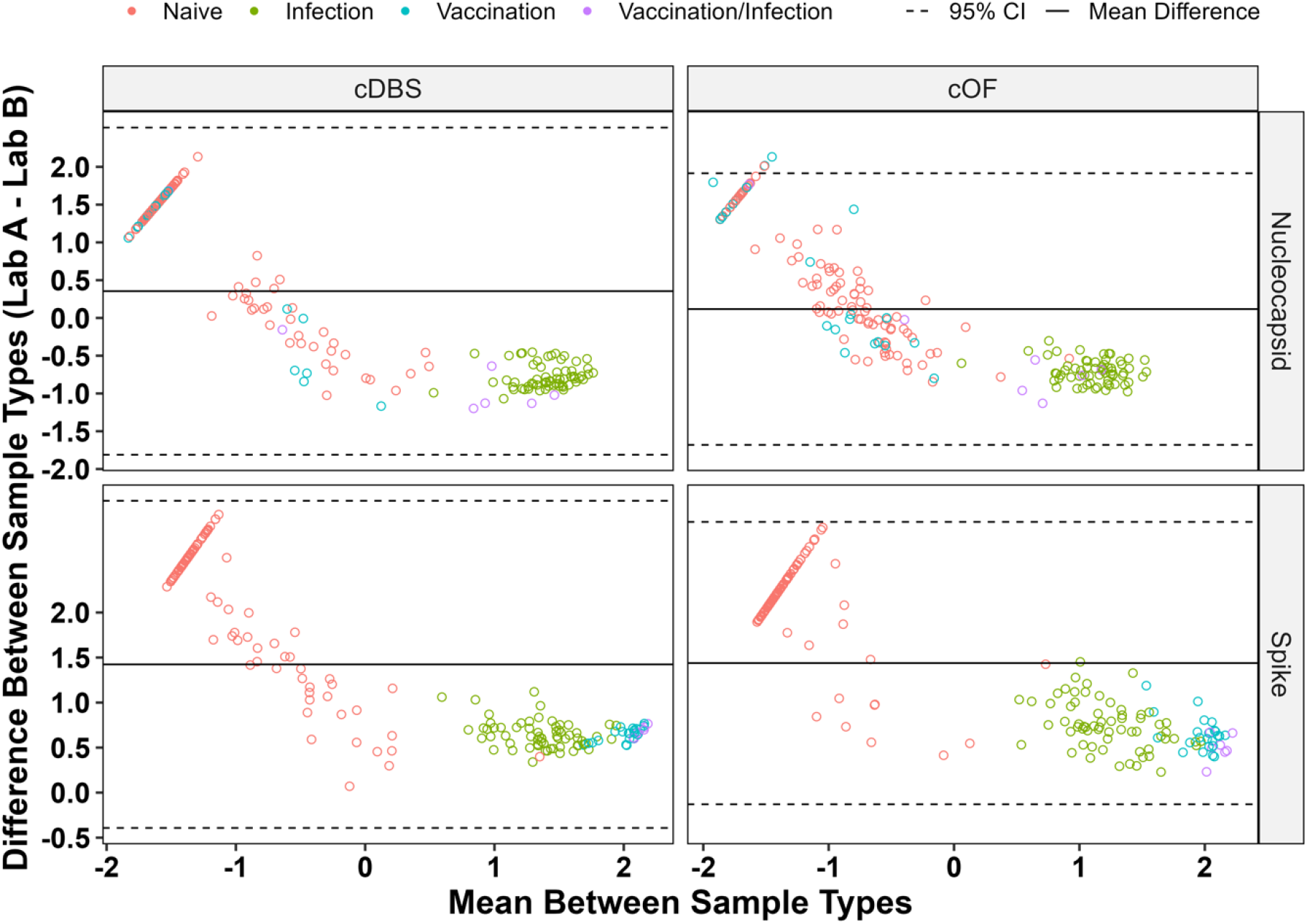
Inter-Laboratory Bland-Altman Analysis for the Lab A and B Assays. Bland Altman plot of 197 paired cDBS and cOF, with comparisons made between labs A and B performing their respective MIAs. Lab A’s MIA uses Mt. Sinai S for cDBS and Sino RBD for cOF while lab B’s MIA uses four spike antigens (RBD, S1, FLS, TRI). Known SARS-CoV-2 antibody status is shown.

### 3.3 : Method Transfer

For the MIA assay platform to be useful within the public health arena, the assays must be transferrable between laboratories. To address the issue of transferability, Lab A adopted Lab B’s assay and Lab B adopted Lab A’s assay using a total of 52 paired OF and DBS samples. When the Lab A assay was performed by labs A and B (Lab A in A/B), results were qualitatively concordant for both cDBS and cOF (κ ≥ 0.92; ≥ 98.1% agreement) (**Table 3**). Kappa coefficients were slightly improved with the lab A MIA, for reasons that are unclear (**Table 3**). Additionally, Bland Altman analysis showed the 52 paired samples were quantitatively concordant (100% within 95% CI) for SARS-CoV-2 reactive cDBS and cOF sampleswith the lab A assay (Lab A in A/B) (**Figure 3, Table S6**)

**Figure 3:**
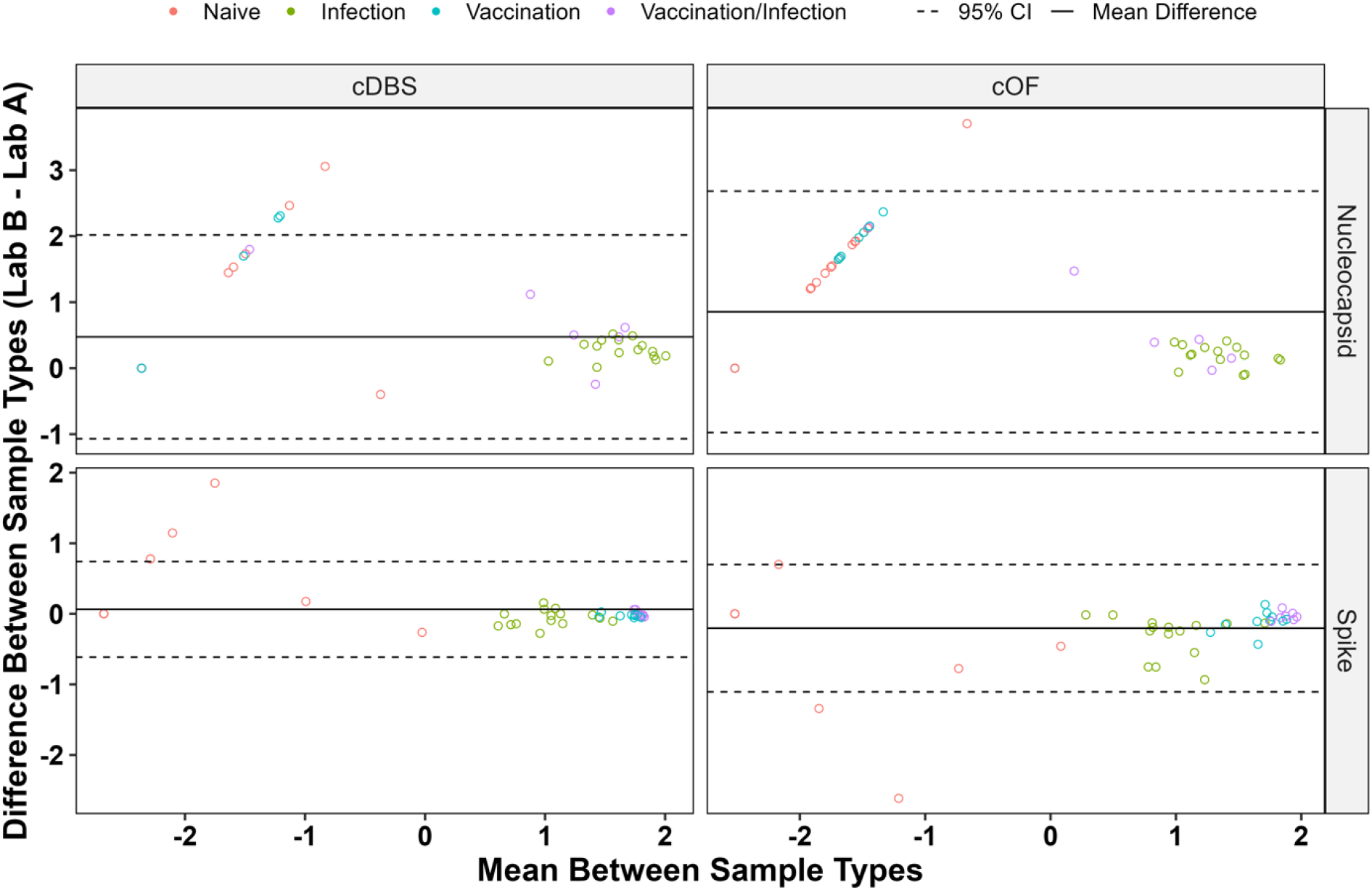
Inter-Laboratory Bland-Altman Analysis for the Lab A Assay. Bland Altman plot of subset of samples (n=52) cDBS and cOF tested by lab A and B using the lab A MIA. Known SARS-CoV-2 antibody status is shown. Inter-laboratory comparison of the lab B assay is shown in supplementary **Figure S2**.

Similarly, when the Lab B assay was performed by labs A and B, (Lab B in A/B), results were qualitatively and quantitively concordant for N (κ = 0.61-1, 82.7%-100% agreement) and S (κ = 0.96-1, 96.2-100% agreement) (**Table 3**). Bland Altman analysis showed the 52 paired samples were quantitatively concordant for the lab B MIA with >96.9% of cDBS and cOF SARS-CoV-2 positive samples falling within the 95% CI (**Figure S2, Table S6**). In conclusion, labs A and B obtained concordant qualitative and quantitative inter-laboratory results for cDBS and cOF when performing method transfer.

## 4. Conclusions and study limitations

In this report we investigated congruity between DBS and OF as model biospecimen types for use in SARS-CoV-2 serological studies. Contrived DBS (cDBS) and OF (cOF) from serum panels from SARS-CoV-2 infected, vaccinated, and uninfected individuals were generated and evaluated using two different SARS-CoV-2 MIAs at two different institutions. With some minor modifications of MIA protocols, we found that DBS and OF were largely interchangeable across laboratories for the qualitative assessment of SARS-CoV-2 antibody determinations. Qualitative assessment was achieved using the same instrument platform (Luminex) but different antigen panels, wash buffers and internal controls, demonstrating that the biospecimens themselves are sufficiently robust to enable interinstitutional comparisons. The obvious limitation of this study is the reliance on contrived biospecimens (DBS and OF), thereby avoiding the inherent variability associated with samples in the field. OF is not a homogenous sample and numerous factors can impact antibody concentration and sample viscosity ^3,9^. Additionally, salivary antibody levels are also known to vary from person to person, and throughout the day for individuals. While DBS are known to be temperature stable for shipping, little is known about OF stability during transportation ^10-12^. Nonetheless, the methods developed in this study are being tested using authentic paired DBS and OF samples to demonstrate intra- and inter-laboratory concordance of these two sample types. Ongoing work will assess IgG and IgA concordance of SARS-CoV-2 antibodies between paired DBS and OF.

## Acknowledgements

We thank Drs. Monica Parker and William Lee of the Wadsworth Center for supervision and technical assistance. We gratefully acknowledge Elizabeth Cavosie for administrative assistance. This study was supported by the National Cancer Institute (NCI) of the National Institutes of Health (NIH) under award number U01CA260508. The content is solely the responsibility of the authors and does not necessarily represent the official views of the NIH.

## Declaration of interest statement

The authors have no conflicts of interest to declare.

**CRediT authorship statement: KLD:** Investigation, Formal analysis, Writing – original draft. **NP:** Investigation, Writing – review & editing. **KK:** Investigation, Writing – review & editing. **LMS:** Conceptualization, Supervision, Writing – review & editing. **MMP:** Conceptualization, Supervision, Writing – review & editing. **CDH:** Conceptualization, Supervision, Writing – review & editing. **NJM:** Conceptualization, Supervision, Funding, Writing – review & editing.

## Notes

### Competing Interest Statement

The authors have declared no competing interest.

