## Supplemental tables and figures for "Inter-laboratory harmonization of microsphere immunoassays for SARS-CoV-2 antibody detection in dried blood spots and oral fluids"

### Supplementary Information

**Table S1. Antigens used in SARS-CoV-2 MIAs.**

| Laboratory | Antigen | Manufacturer | Catalog # |
|---|---|---|---|
| Lab A | Gen N – Nucleoprotein | GenScript | Z03480 |
| | NAC – Nucleoprotein | Native Antigen | REC31851 |
| | Sino RBD – RBD | Sino Biological | 40592-V08H |
| | Mt. Sin RBD – RBD | Mt. Sinai | |
| | Gen RBD – RBD | GenScript | Z03483 |
| | Mt. Sin S – Spike | Mt. Sinai | |
| | ECD – S1+S2 ectodomain | Sino Biological | 40589-V08B1 |
| | SARS-CoV Nucleoprotein | Native Antigen | REC31744 |
| | hCoV 229E S1+S2 ectodomain | Sino Biological | 40605-V08B |
| | hCoV OC43 hemagglutinin esterase | Sino Biological | 40603-V08H |
| | hCoV HKU1 isolate N1 S1 | Sino Biological | 40021-V08H |
| | hCoV NL63 S1+S2 ectodomain | Sino Biological | 40604-Vo8B |
| | SARS-CoV RBD | Sino Biological | 40150-V08B2 |
| | MERS-CoV S1 | Native Antigen | REC31760 |
| | RSV A2 | Sino Biological | 10049-V08B |
| | RSV RSS2 | Sino Biological | 40037-V08B |
| | Control anti-IgG | Jackson Immunolabs | 109-005-098 |
| | Control anti-IgA | Jackson Immunolabs | 109-005-011 |
| | Control anti-IgM | Jackson Immunolabs | 109-005-129 |
| | Control BSA | Luminex | 30-00136 |
| Lab B | N NA - Nucleocapsid | Native Antigen | REC31851 |
| | N-SB – Nucleocapsid | Sino Biological | 40588-V08B |
| | NHT - Nucleocapsid | Sino Biological | 40588-V07E |
| | RBD – Spike RBD | Mass Biologics | |
| | S1 – Spike S1 | Sino Biological | 40591-V08H |
| | FLS – Full length spike | Native Antigen | REC31868 |
| | TRI – Spike trimer | Mass Biologics | |
| | IgG3 internal control | ThermoFisher | MA183242 |
| | Control BSA | Luminex | 30-00136 |

**Table S2: Reactive cutoff MFIs for the SARS-CoV-2 MIAs.**

| MIA | Antigen | cDBS Cutoff | Serum Cutoff | cOF Cutoff |
|---|---|---|---|---|
| Lab A | Gen N | 230 | 230 | 330 |
| | NAC N | 2,000 | 2000 | 330 |
| | Sino RBD | 340 | 340 | 330 |
| | Mt. Sin S | 475 | 475 | 500 |
| Lab B | N NA | 761 | 3,766 | 1,521 |
| | RBD | 1,066 | 2,660 | 1,255 |
| | S1 | 295 | 215 | 170 |
| | FLS | 330 | 469 | 361 |
| | TRI | 589 | 2,801 | 1,249 |

Table S1. Commercially available serum panels used for 197 paired contrived specimens (100 SARS-CoV-2 negative, 97 SARs-CoV-2 positive).

| **Panel** | **n** | **Manufacturer** | **Infection/Vaccination Status** |
|---|---|---|---|
| SARS-CoV-2 convalescent plasma | 3 | Access Biologicals | Infection |
| SARS-CoV-2 seroconversion | 7 | Access Biologicals | Infection |
| SARS-CoV-2 negative (Panel E) | 77 | Access Biologicals | Naïve |
| SARS-CoV-2 IgG and IgM positive (Panel D) | 30 | Access Biologicals | Infection |
| SARS-CoV-2 IgG positive (Panel F) | 20 | Access Biologicals | Infection |
| SARS-CoV-2 post-vaccine series (Panel H) | 23 | Access Biologicals | Vaccination |
| SARS-CoV-2 post-vaccine series (Panel H) | 7 | Access Biologicals | Infection/vaccination |
| SARS-CoV-2 pre-vaccine series (Panel H) | 23 | Access Biologicals | Naïve |
| SARS-CoV-2 pre-vaccine series (Panel H) | 6 | Access Biologicals | Infection |
| SARS-CoV-2 Delta positive | 1 | WC | Infection |

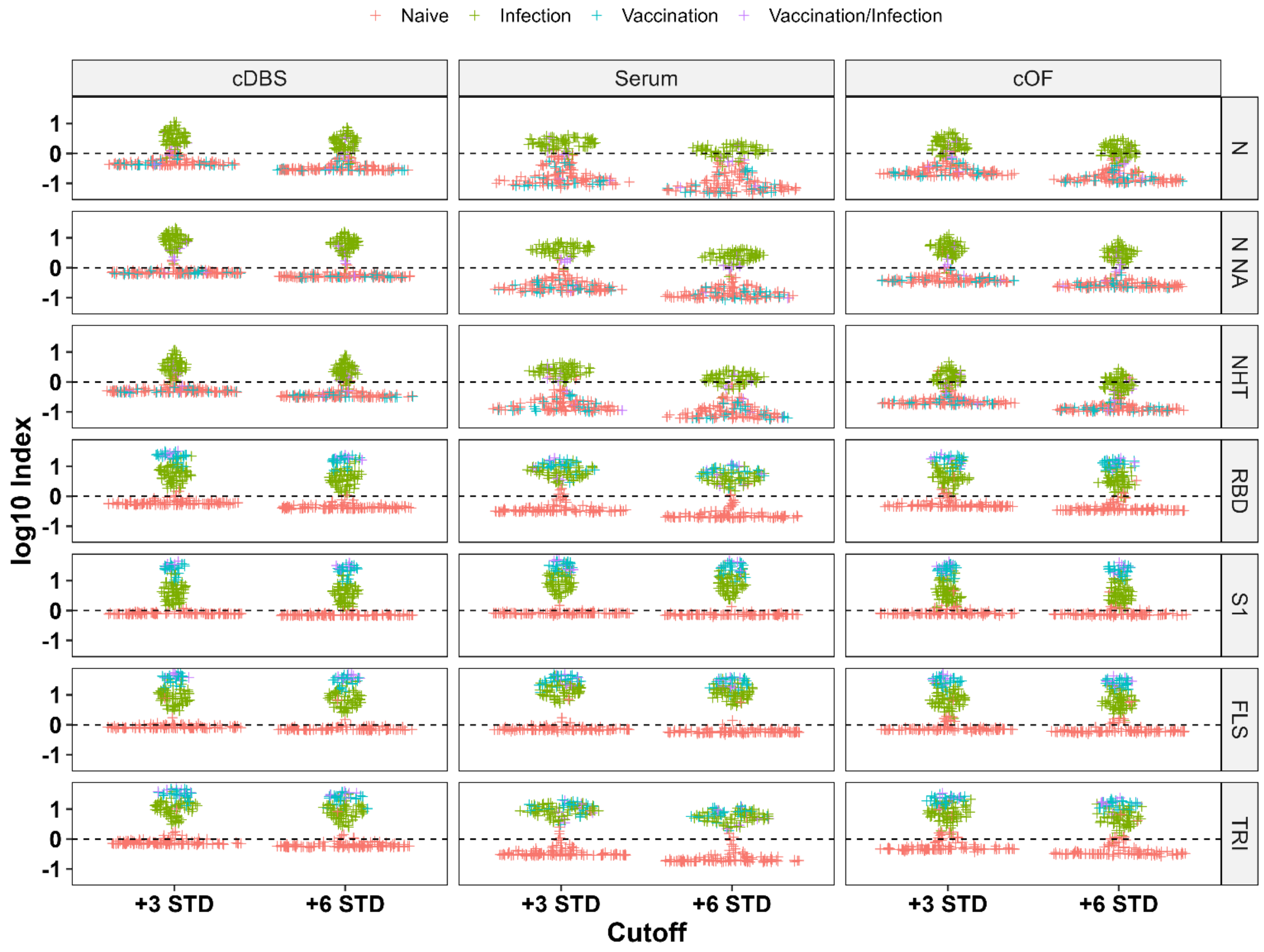


Figure S1: Comparison of +3 STD +6 STD cutoff values in A) cDBS samples, B) serum samples, and C) cOF samples for the lab B assay. Known SARS-CoV-2 antibody status is shown.

Table S4: Percentage of SARS-CoV-2 positive (N = 74; S = 97) and negative (N = 123; S = 100) samples within the 95% CI for paired cDBS and cOF Bland Altman plots shown in Figure 1. Concordance is defined as ≥ 95% of data points falling within the 95% CI.

| **Laboratory** | **Positive (%)** | | **Negative (%)** | |
|---|---|---|---|---|
| | **Nucleocapsid** | **Spike** | **Nucleocapsid** | **Spike** |
| Lab A | 100.0 | 97.9 | 94.3 | 92.0 |
| Lab B | 100.0 | 100.0 | 93.5 | 84.0 |

Table S5: Coefficient of variation for 12 technical replicates of positive controls for cDBS, serum, and cOF under various assay and wash buffer conditions for the lab B MIA (assay buffer/wash buffer). Assay and wash buffer either contain (+) or lack (-) BSA. Buffers detailed below.

| **Sample Type** | **Buffer** | **N NA (%)** | **RBD (%)** | **S1 (%)** | **FLS (%)** | **TRI (%)** |
|---|---|---|---|---|---|---|
| cDBS | *+/+ | 7.0 | 5.3 | 8.7 | 5.9 | 7.9 |
| | -/+ | 64.1 | 56.0 | 65.8 | 60.7 | 57.6 |
| | -/- | 10.4 | 14.0 | 12.5 | 9.6 | 9.7 |
| Serum | +/+ | 35.4 | 27.3 | 31.7 | 24.9 | 20.1 |
| | -/+ | 43.7 | 32.2 | 43.1 | 37.1 | 31.8 |
| | *-/- | 19.1 | 19.4 | 22.7 | 17.4 | 16.0 |
| cOF | +/+ | 83.7 | 99.2 | 94.4 | 90.9 | 80.6 |
| | -/+ | 76.7 | 71.5 | 90.3 | 71.1 | 63.4 |
| | *-/- | 15.1 | 14.6 | 17.5 | 16.4 | 15.1 |

**Optimal buffer conditions; Assay Buffer +: PBS + 2% BSA; Assay Buffer -: PBS-TBN; Wash Buffer +: PBS + 2% BSA + 0.02% Tween 20 + 0.05% sodium azide; Wash Buffer -: PBS + 0.05% Tween20*

Table S6: Percentage of SARS-CoV-2 positive (N = 74; S = 97) and negative (N = 123; S = 100) samples within the 95% CI for paired cDBS and cOF Bland Altman plots for inter-laboratory comparisons (n = 197) between labs A and B (Figure 2). Also shown is the percentage of SARS-Cov-2 positive (N = 22; S = 32) and negative (N = 30; S = 20) for method transfer comparisons between labs A and B (n = 52) (Figure 3). Concordance is indicated by ≥ 95% of data points falling within the 95% CI.

| **MIA Performed** | **Sample Type** | **Positive** | | **Negative** | |
|---|---|---|---|---|---|
| | | **Nucleocapsid** | **Spike** | **Nucleocapsid** | **Spike** |
| **Lab A and B, respectively** | cDBS | 100.0 | 100.00 | 100.00 | 100.0 |
| | cOF | 100.00 | 100.0 | 97.6 | 100.0 |
| **Lab A** | cDBS | 100.0 | 100.0 | 86.7 | 85.0 |
| | cOF | 100.0 | 100.0 | 96.7 | 85.0 |
| **Lab B** | cDBS | 100.0 | 100.0 | 90.0 | 85.0 |
| | cOF | 100.0 | 96.9 | 90.0 | 100.0 |

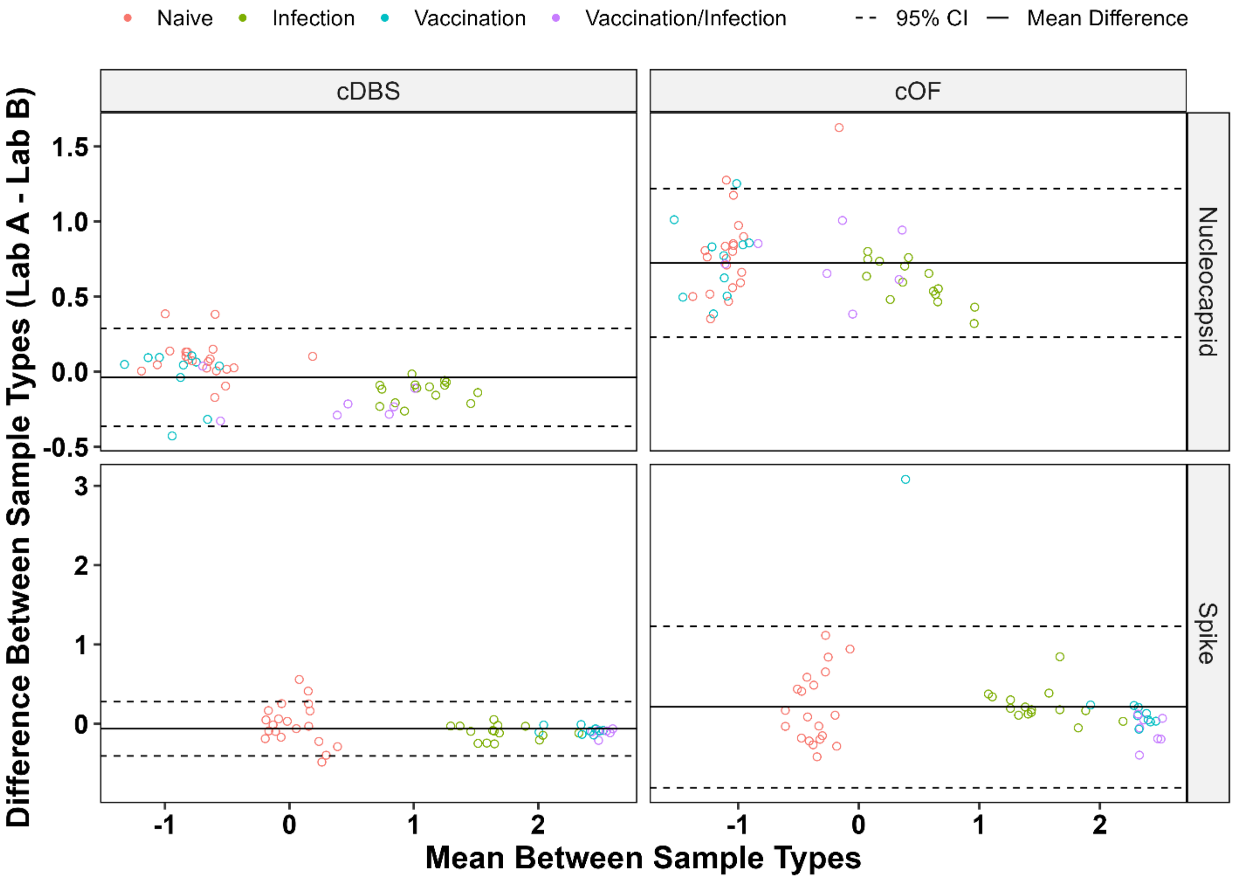


Figure S2: Bland Altman plot of subset of samples (n=52) cDBS and cOF tested by lab A and B using the lab B MIA. Known SARS-CoV-2 antibody status is shown.
